# DPAC: a tool for Differential Poly(A) Site usage from poly(A)–targeted RNAseq data

**DOI:** 10.1101/531590

**Authors:** Andrew Routh

## Abstract

Poly(A)-tail targeted RNAseq approaches, such as 3’READS, PAS-seq and Poly(A)-ClickSeq, are becoming popular alternatives to random-primed RNAseq for simplified gene expression analyses as well as to measure changes in poly(A) site usage. We and others have recently demonstrated that these approaches perform similarly to other RNAseq strategies, while saving on the volume of sequencing data required and providing a simpler library synthesis strategy. Here, we present DPAC; a streamlined pipeline for the preprocessing of poly(A)-tail targeted RNAseq data, mapping of poly(A)-sites and poly(A) clustering, and determination of differential poly(A) site usage using DESeq2. Changes in poly(A) site usage is simultaneously used to report differential gene expression, differential terminal exon usage and alternative polyadenylation (APA).

## Introduction

The abundance of RNA transcripts as well as poly(A) site positions can be determined directly from a number of RNAseq techniques that focus reads to the junction of 3’UTRs and poly(A) tails. Numerous approaches, including as 3’READS [1], PAS-seq [2] and Poly(A)-ClickSeq [3] are commonly and commercially available and can be used to estimate transcript abundances, differential gene expressions, alternative terminal exon usage (TE) and alternative poly-adenylation (APA). In addition to providing information on the location of poly(A)-sites (PASs) in mRNA transcripts, these methods provide a simple and equally powerful alternative to randomly-primed RNAseq in both bulk and single-cell RNAseq experiments. We and others have recently demonstrated that poly(A)-targeted RNAseq approaches perform differential expression analyses similarly to other RNAseq strategies, while saving on the volume of sequencing data required and providing a simpler library synthesis strategy [4].

We present DPAC as a pipeline to preprocess raw poly(A)-tail target RNAseq data, determine the location of PASs and determine the differential abundance of PASs between two conditions. DPAC comprises three major stages; 1) Pre-processing of raw poly(A)-tailed RNAseq including estimation of poly(A)-tail lengths and mapping to a reference genome; 2) an optional step that locates all PASs in the provided data and generates annotated poly(A) clusters (PACs); and 3) a differential expression analysis of PACs using DESeq2. In addition to determining changes in individual PAS abundance, DPAC will also calculate changes in terminal exon usage and gene expression by collapsing read counts form individual PASs if they are present on the same exon/intron and whole-gene respectively. DPAC compiles these results and generates a final output table simultaneously describing changes in gene expression, terminal exon (or intron) usage and alternative poly-adenylation.

We demonstrate the utility of this pipeline by re-analyzing our previously published data using Poly(A)-ClickSeq to measure changes in PAS usage in HeLa cells knocked-down for mammalian Cleavage Factor I 25kDa subunit (CF25Im) [3]. As expected, DPAC reports that CF25Im depletion results in substantial shortening in 3’UTRs, while only minimally affecting overall gene expression levels. DPAC, along with annotated PAS cluster databases, is maintained and available at https://sourceforge.net/projects/DPAC/

## Methods

DPAC is a simple bash batch script with associated python scripts, run with a single command line entry. The pipeline can be broken down into 4 main stages, each of which can be invoked independently to allow re-analysis with new parameters. A number of software dependencies are listed, though these are all common in RNAseq pipelines and on bioinformatic servers. A flow chart of each of the main stages of DPAC is shown in **Figure 1**.

**Figure 1:**
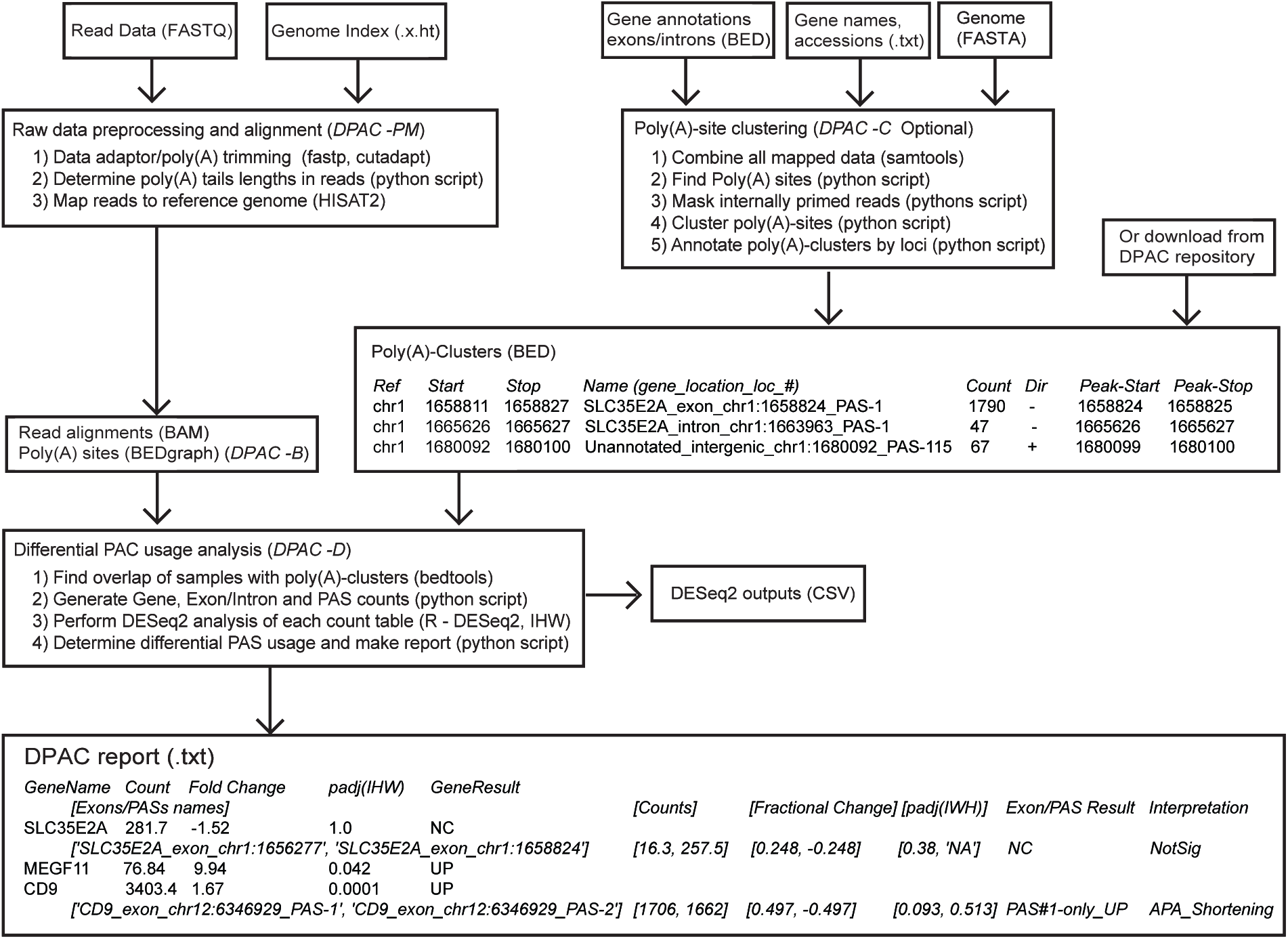
A flow-chart summarizing each of the stages, required input files and returned output files for the DPAC pipeline. Command-line options used to invoke each stage are illustrated: DPAC –PM for preprocessing and Mapping, -C for poly(A) cluster generation, -B for bedgraphs, -D for the final differential PAC usage analysis. Examples of the output of the DPAC pipeline are shown for three genes: SCL35E2A, MEGF11, and CD9.

### 2.1 Initial Data Prep and poly(A) site mapping

3’ end sequencing methods including Poly(A)-ClickSeq (PAC-Seq) [3]generate raw sequence reads overlapping the junction of the 3’ UTR and the poly(A) tail of mRNA transcripts. The preparation of raw read data in terms of adapter trimming, poly(A) tail trimming, poly(A) tail length and quality filtering are the same as previously described in Routh et al. [3]. Mapping to a reference genome as well as extraction of poly(A) events are also performed as previously described. Briefly, reads are trimmed and quality filtered using *fastp* [5] (parameters: -a AGATCGGAAGAGC -f 6 -g -l 40 –Q). Trimmed reads are trimmed a second time using *cutadapt* [6] to remove and measure the poly(A) tail (parameters: -b A{15} -n 2 -O 6 -m 40). Using the script (Extract_pA_Lens.py), reads from output from this step are compared to the input to determine how many A’s (if any) were removed from the 3’ end of the read. This number is appended to the name of each read for future quality filtering. The preprocessing steps of DPAC are invoked by default or by using the ‘–p P’ command-line argument.

After data preparation, reads are mapped using default settings to the reference genome using *HiSat2 [7]*. Again, the mapping step of DPAC is invoked by default or by using the ‘–p M’ command-line argument. If required, DPAC will also output the individual bedgraphs illustrating all poly(A)-sites and mapping coverage for each sample when using the command-line argument ‘–p B’. These files can be loaded into canonical genome browsers and may be useful when generating figures. An example of these output data is shown in **Figure 2,** illustrating the mapping of PAC-Seq reads and identified PASs for two samples. However, they are not required for the downstream analysis.

**Figure 2:**
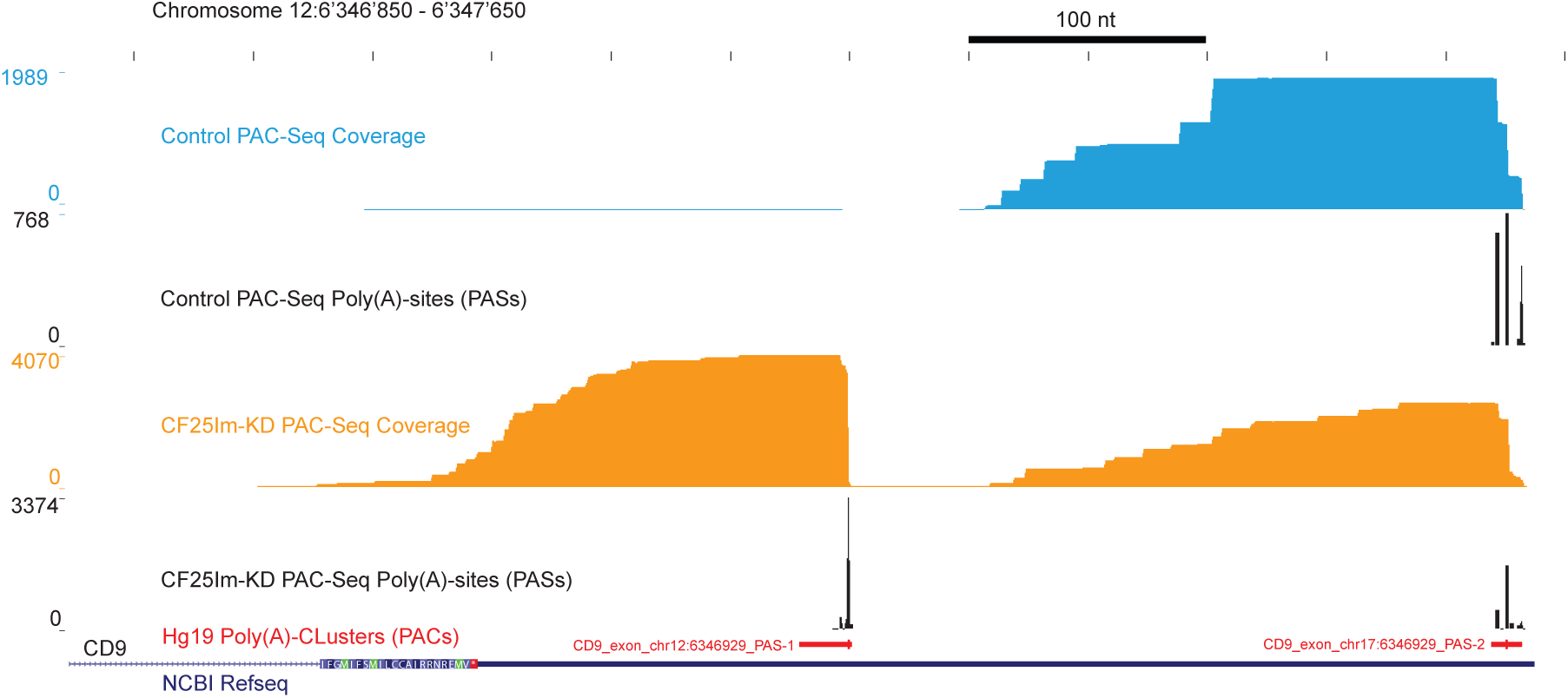
Read coverage and the detected poly(A) sites over the CD9 gene for two samples of Poly(A)-ClickSeq analysis of mocked treated HeLa cells (blue) and CF25Im siRNA treated HeLa cells (orange) are depicted. Poly(A)-Clusters are illustrated as a track (red) in the UCSC genome browser. The most frequently detected poly(A)-site within the poly(A) cluster is highlighted as the thicker portion of the whole poly(A) cluster in the track.

### 2.2 Making poly(A) clusters

To maximize the power of PAS annotation using PAC-Seq, the data from all samples is first combined and the 3’end of all reads are used to identify the location of PASs across the reference genome. The number of non-primer derived As found in each read is utilized to score the confidence of the PAS. By default, PASs are outputted to a raw BEDGraph file if a poly(A) site is identified by at least 5 reads with at least 5 extra non-primed-derived A’s per read. Finally, PASs are filtered for internal priming by counting the number of A’s in the reference genome immediately downstream of the identified PASs. If 12 or more As are found within 20 nts downstream, these events are ‘masked’ and not further utilized.

PASs are predominantly found at a ‘GA’, ‘UA’ or ‘CA’ dinucleotides, although the exact site is variable [8, 9]. Single PASs occurring within 25nts of one another are merged into poly(A)-clusters (PACs), which are subsequently treated as singular features in downstream analysis. Information regarding the most frequent PAS nucleotide is retained as extra columns in the output BED file. To annotate the new PACs, locations of exonic and intronic sequences are obtained UCSC database. The overlap of each PAC to annotated exons and introns is then determined using *bedtools* [10]. PACs are named according the gene name, whether the PAC is exonic, intronic or found just downstream of a terminal exon, followed by the genomic coordinate, and then assigned a number depending upon the number of other PACs found within the same exon or intron. PACs found in intergenic or otherwise unannotated sequences are numbered sequentially depending number of unannotated PACs found. This naming scheme is used in the final stage of DPAC to differentiate between alternative poly-adenylation events and alternative terminal exon usage.

The poly(A) clustering stage of DPAC requires specific BED files of annotated genes, exons and introns which can be obtained from the UCSC genome browser table browser. This is not invoked by default, but by using the command-line entry ‘-p C’. Pre-compiled databases of PACs generated using large PAC-Seq sequencing datasets, such as the ones used in this report, can be found online at https://sourceforge.net/projects/dpac-seq/files/Poly%28A%29_Clusters_BED/ These can be downloaded and used for the final stage of DPAC. Examples of two specific poly(A) clusters within the CD0 gene are illustrated in **Figure 2** in the hg19 Poly(A)-Cluster track in red.

### 2.3 Determination of differential exon usage using DESeq2

In the final stage of DPAC, the mapped reads from each individual samples are used to determine the frequency of PAC in each dataset by determining the overlaps of the 3’ ends of the mapped reads with the provided poly(A) cluster database using *bedtools* [10]. A PACs is counted if the 3’ end of a mapped read overlaps within a user-defined distance (10nts is default value) of an annotated poly(A) cluster and count tables of PACs are returned. If multiple PASs are found within an exon or intron, then these are collapsed into a single entry, generating a new count table for exons and introns. Similarly, if multiple PASs are found within a single gene, these are also collapsed to create a count table just for whole genes. However, by default, only PASs found in exonic regions are collapsed into gene counts as introns can often contain repetitive and/or transposable element whose inclusion can artificially inflate count numbers (however, this parameter can be overturned to force inclusion of intronic PASs). Three sets of count tables are thus generated (see example for CD9 in **Table 1**) and passed individually into DESeq2 [11]. Data normalization and statistical tests are applied using the canonical DESeq2 pipeline using *local* dispersion estimation and Independent Hypothesis Weighting (IHW) [12] to estimate false discovery rates and power maximization. Thus, differential usage of PASs, exons/introns and whole-genes are each calculated and the results are output as csv files. As illustrated in the flow-chart in **Figure 1**, these files are returned for inspection, figure generation and other downstream analyses.

**Table 1:**
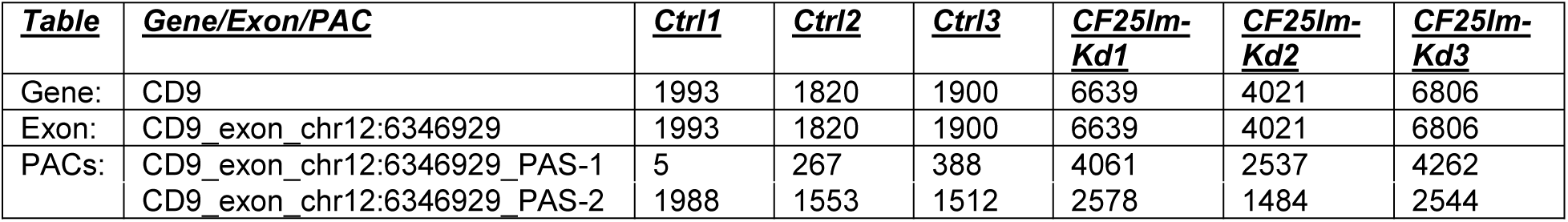
Example of count table used or DESeq2 for CD9, CD9 exon, and CD9 poly(A) clusters.

### 2.4 Output

After DESeq2 analysis, a final compiled table is generating containing information about PAS usage for each gene. If a gene only has one PASs site and thus one terminal exon, only the gene information is returned. Genes with differential expression (fold-change > 1.5; padj < 0.1) are annotated as ‘DOWN’ or ‘UP’. If there is no significant change, genes are labels as ‘NC’ (No Change). Alternative polyadenylation (APA) or differential terminal exon usage is reported when a gene has two or more PASs (minimum occupancy of 5% per PAS), with at least one PAS undergoing differential usage with an IHW padj <0.1 and resulting in a fractional change of the PAS usage by at least 10%. If the two PASs are found in the same exon, then this annotated as an APA event. The locations of the two (or more) PASs are then used to determine whether the APA results in 3’UTR shortening or lengthening. If two PASs are found in different exons, this is annotated as a terminal exon usage event (denoted as ‘TE’).

### 2.5 Data Availability Statement

Data used in the manuscript is available at the NCBI SRA database with project number *PRJNA374982*. DPAC is freely available (MIT license), along with annotated PAS cluster databases, is maintained and available at https://sourceforge.net/projects/DPAC/

## Results and Discussion

To evaluate this pipeline, we re-analyzed the PAC-seq data deposited at NCBI SRA (*PRJNA374982*) from the original PAC-Seq publication [3]. Six datasets derived from total cellular RNA extracted from three technical replicates each of mock-treated and CF25Im KD HeLa cells. We applied our pipeline to locate and annotate *de novo* poly(A) clusters and then to determine the differential usage of PACs between each condition. Annotation data and example command-line entries are provided in the DPAC manual to repeat these analyses.

During *de novo* PAS clustering a total of 45’587 poly(A) clusters (PACs) with >30 reads were identified in the datasets. Of these, 28’146 were exonic, 7’950 were intronic, 1’168 were found within 250nts downstream of annotated 3’ terminal exons; and 8’314 were intergenic or otherwise unannotated (**Supplementary Data 1**).

Our pipeline found PACs mapped over a total of 13’778 genes, 1525 of which exhibited multiple terminal exons and 7178 exhibited multiple PASs (52%, similar to previously reported [3, 13]). From these events, DPAC reported differential expression of 378 genes (**Supplemental Data 2**). We found that 1134 individual PACs were differentially expressed (IHW-padj < 0.1, fold-change >1.5) (**Supplemental Data 3**). Volcano plots illustrating these changes in gene expression and PAC abundance are shown in **Figure 3**. Due to changes in PAC abundance, DPAC reported that 621 genes exhibited APA with the 3’UTR shortening of 452 exons and lengthening of 66 exons. 103 exons exhibited both lengthening and shortening due to the presence of multiple PASs within single exons. Differential PAS usage resulted in the alternative terminal exon usage in only 44 genes and only 9 genes exhibited APA and TE simultaneously. The entire report is available in **Supplemental Data 4**.

**Figure 3:**
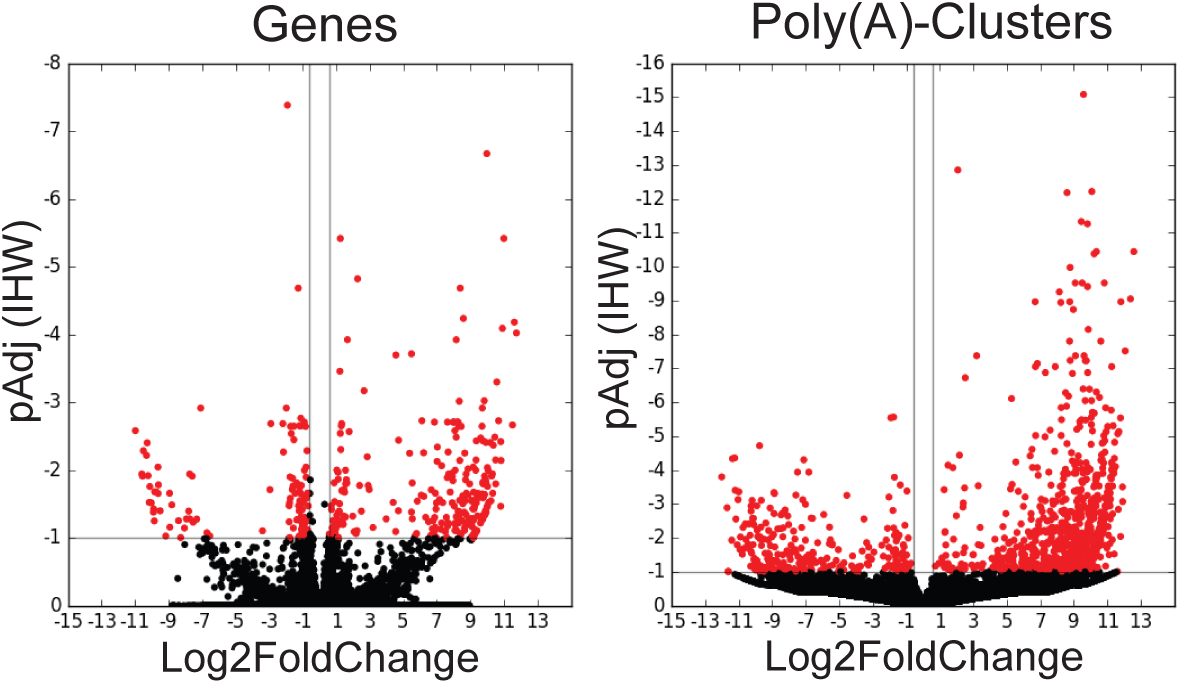
Volcano plots of the differential expression of Genes (left) and Poly(A)-Clusters (right) as determined by DESeq2. Red dots indicate genes or PACs with a fold changes greater than 1.5 and a p-adjusted value less than 0.1.

Specific examples of the final output are shown in **Figure 1**. SLC35E21 has two PASs found in two different terminal exons, but there is no significant change in their usage and therefore no alternative terminal exon usage. MEGF11 has only one PAS, and this gene significantly up-regulated upon CF25Im KD. CD9 is also up-regulated upon CF25Im KD due to the up-regulation of one of two PACs found within the same terminal exon with a net effect of 3’UTR shortening. The mapping of the raw data over CD9 and the detected PASs are shown in **Figure 2**.

The predominant shortening of 3’UTRs upon knock-down of CF25Im is the expected phenotype and is consistent with our and other previous analyses [3, 14]. By virtue of measuring differential usage of each individual PAS independently, this pipeline revealed DE of PASs outside of annotated regions. Indeed, of the 1134 identified DE APA sites, 113 (10.0%) were intronic and 109 (9.6%) were found in unannotated regions. Interestingly, by comparing to *repeatmasker* database [15], 80 of these PACs mapped next to repetitive and/or transposable elements (e.g. *Alu* elements), indicating that these elements may also be differentially regulated upon the modulation of CF25Im activity.

In summary, DPAC performs each of the necessary steps required for preprocessing, PASs identification and differential PAS usage required for poly(A)-targeted RNAseq experiments. The pipeline reports the expected findings upon reanalysis of Poly(A)-ClickSeq datasets comparing mock and CF25Im-knockdown HeLa cells. By virtue of assessing changes in all PAS regardless of whether they are found in annotated genomic regions, DPAC may also allow discovery of novel mRNA transcripts and/or changes in the expression of ncRNAs and/or non-coding transposable elements. DPAC therefore provides a singular pipeline to simultaneously report differential gene expression, terminal exon usage and alternative poly-adenylation.

## Supporting information

DPAC software; Scripts, Manual and TestData

## Acknowledgements

A.R. is supported by start-up funds from the University of Texas Medical Branch. I thank Dr. Eric Wagner and Dr. Mugé Martinez-Kuyumcu at UTMB for providing critical support and advice.

