## Supplementary material for "DPAC: a tool for Differential Poly(A) Site usage from poly(A)–targeted RNAseq data": DPAC software; Scripts, Manual and TestData: DPAC_Manuscript.pdf

Andrew Routh[1,2]

1) Department of Biochemistry and Molecular Biology, The University of Texas Medical Branch, Galveston, TX, USA
2) Sealy Centre for Structural Biology and Molecular Biophysics, University of Texas Medical Branch, Galveston, Texas, USA.

**Table 1**: Example of count table used or DESeq2 for CD9, CD9 exon, and CD9 poly(A) clusters.

| ***Table*** | ***Gene/Exon/PAC*** | ***Ctrl1*** | ***Ctrl2*** | ***Ctrl3*** | ***CF25Im-Kd1*** | ***CF25Im-Kd2*** | ***CF25Im-Kd3*** |
|---|---|---|---|---|---|---|---|
| Gene: | CD9 | 1993 | 1820 | 1900 | 6639 | 4021 | 6806 |
| Exon: | CD9_exon_chr12:6346929 | 1993 | 1820 | 1900 | 6639 | 4021 | 6806 |
| PACs: | CD9_exon_chr12:6346929_PAS-1 | 5 | 267 | 388 | 4061 | 2537 | 4262 |
| | CD9_exon_chr12:6346929_PAS-2 | 1988 | 1553 | 1512 | 2578 | 1484 | 2544 |

A.R. is a co-founder and owner of ‘ClickSeq Technologies LLC’, which is a Texas-based Next-Generation Sequencing solution provider offering ClickSeq protocols including PAC-Seq.

### ***Figure Legends***

**Figure 1**: A flow-chart summarizing each of the stages, required input files and returned output files for the DPAC pipeline. Command-line options used to invoke each stage are illustrated: DPAC –PM for preprocessing and Mapping, -C for poly(A) cluster generation, -B for bedgraphs, -D for the final differential PAC usage analysis. Examples of the output of the DPAC pipeline are shown for three genes: SCL35E2A, MEGF11, and CD9.

**Figure 2**: Read coverage and the detected poly(A) sites over the CD9 gene for two samples of Poly(A)-ClickSeq analysis of mocked treated HeLa cells (blue) and CF25Im siRNA treated HeLa cells (orange) are depicted. Poly(A)-Clusters are illustrated as a track (red) in the UCSC genome browser. The most frequently detected poly(A)-site within the poly(A) cluster is highlighted as the thicker portion of the whole poly(A) cluster in the track.

**Figure 3**: Volcano plots of the differential expression of Genes (left) and Poly(A)-Clusters (right) as determined by DESeq2. Red dots indicate genes or PACs with a fold changes greater than 1.5 and a p-adjusted value less than 0.1.

### ***References***


[1] Zheng D, Liu X, Tian B. 3'READS+, a sensitive and accurate method for 3' end sequencing of polyadenylated RNA. RNA. 2016;22:1631-9.
[2] Shepard PJ, Choi E-A, Lu J, Flanagan LA, Hertel KJ, Shi Y. Complex and dynamic landscape of RNA polyadenylation revealed by PAS-Seq. RNA. 2011;17:761-72.
[3] Routh A, Ji P, Jaworski E, Xia Z, Li W, Wagner EJ. Poly(A)-ClickSeq: click-chemistry for next-generation 3-end sequencing without RNA enrichment or fragmentation. Nucleic Acids Res. 2017;45:e112.
[4] Elrod NR, Jaworski EA, Ji P, Wagner EJ, Routh A. Development of Poly(A)-ClickSeq as a Tool Enabling Simultaneous Genome-wide Poly(A)-site identification and Differential Expression Analysis. Methods. 2019.
[5] Chen S, Zhou Y, Chen Y, Gu J. fastp: an ultra-fast all-in-one FASTQ preprocessor. Bioinformatics. 2018;34:i884-i90.
[6] Martin M. Cutadapt removes adapter sequences from high-throughput sequencing reads. EMBnetjournal. 2011;17:10-2.
[7] Kim D, Langmead B, Salzberg SL. HISAT: a fast spliced aligner with low memory requirements. Nature methods. 2015;12:357-60.
[8] Routh A, Ji P, Jaworski E, Xia Z, Li W, Wagner EJ. Poly(A)-ClickSeq: click-chemistry for next-generation 3-end sequencing without RNA enrichment or fragmentation. Nucleic Acids Res. 2017.
[9] Derti A, Garrett-Engele P, Macisaac KD, Stevens RC, Sriram S, Chen R, et al. A quantitative atlas of polyadenylation in five mammals. Genome Res. 2012;22:1173-83.
[10] Quinlan AR, Hall IM. BEDTools: a flexible suite of utilities for comparing genomic features. Bioinformatics. 2010;26:841-2.
[11] Love MI, Huber W, Anders S. Moderated estimation of fold change and dispersion for RNA-seq data with DESeq2. Genome Biol. 2014;15:550.
[12] Ignatiadis N, Klaus B, Zaugg JB, Huber W. Data-driven hypothesis weighting increases detection power in genome-scale multiple testing. Nature methods. 2016;13:577-80.
[13] Lee JY, Yeh I, Park JY, Tian B. PolyA_DB 2: mRNA polyadenylation sites in vertebrate genes. Nucleic Acids Res. 2007;35:D165-8.
[14] Zhu Y, Wang X, Forouzmand E, Jeong J, Qiao F, Sowd GA, et al. Molecular Mechanisms for CFIm-Mediated Regulation of mRNA Alternative Polyadenylation. Mol Cell. 2018;69:62-74 e4.
[15] Smit A, Hubley, R & Green, P. RepeatMasker Open-4.0. http://wwwrepeatmaskerorg. 2013-2015.

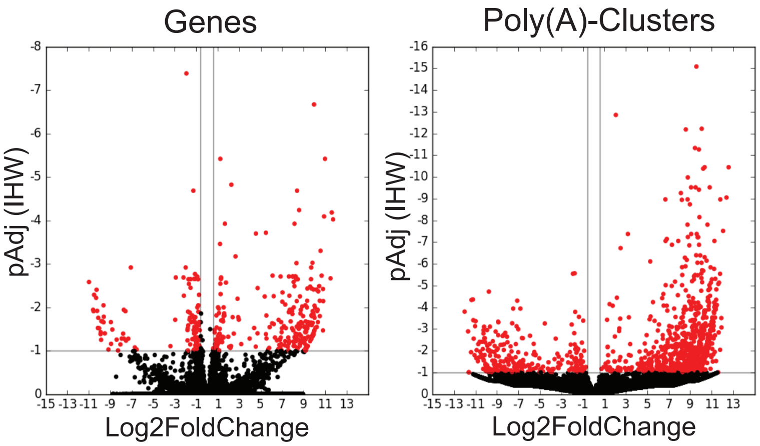


Figure 3 - Volcano plots, Genes and PACs

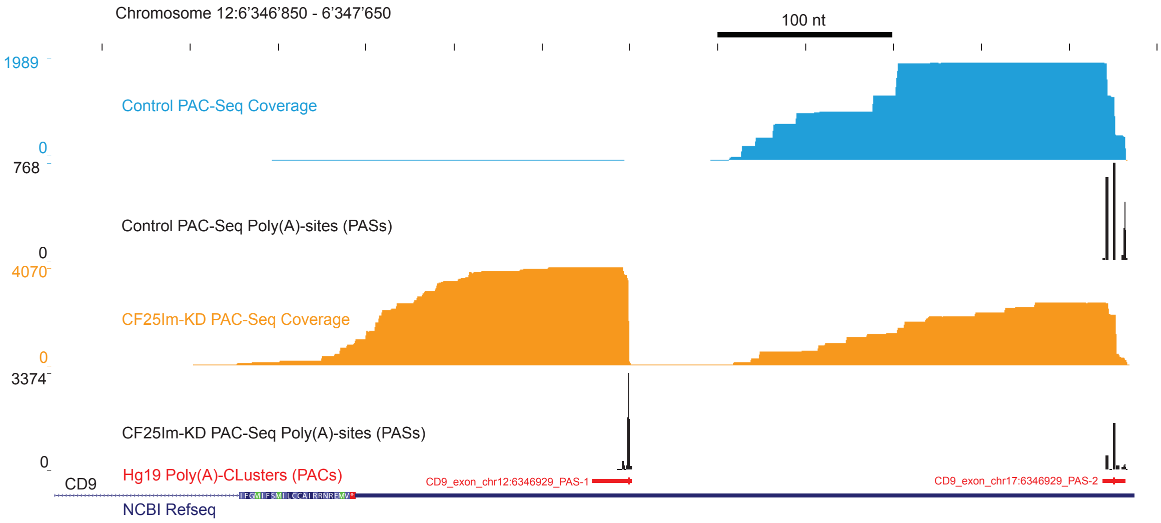


Figure 2 - Mapping visualisation in UCSC Genome Browser

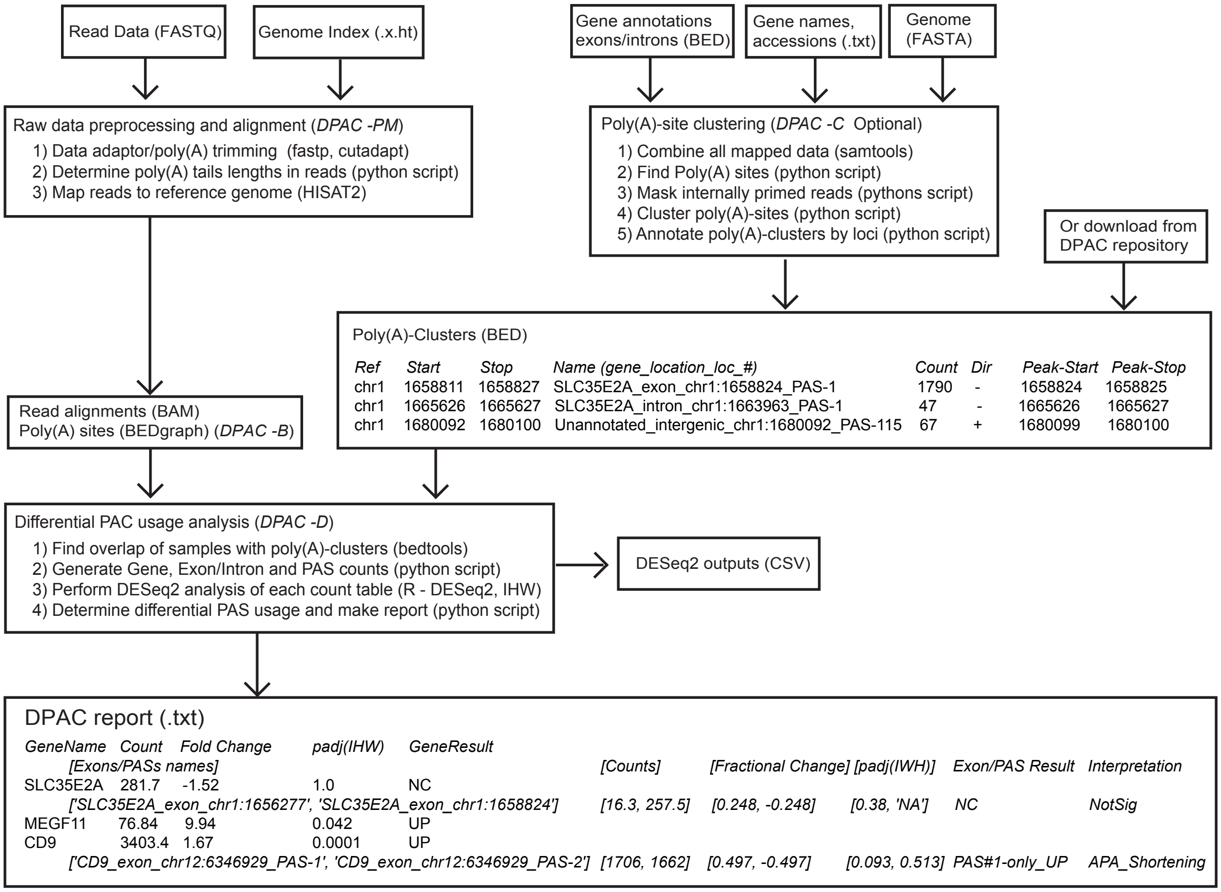


Figure 1 - DPAC Flow Chart
