## Supplementary material for "DPAC: a tool for Differential Poly(A) Site usage from poly(A)–targeted RNAseq data": DPAC software; Scripts, Manual and TestData: APA_Terminal_Exon.py

```
#!/bin/python
import argparse
import numpy as np
np.seterr(divide='ignore', invalid='ignore')

parser = argparse.ArgumentParser()
parser.add_argument("Root", help="Root of Datafiles")
parser.add_argument("Output", help="Report Name")
parser.add_argument("PASs", help="Report Name")
parser.add_argument("--Replicates", help="Report Name")
parser.add_argument("--Conditions", help="Report Name")
parser.add_argument("--Fraction", help="Report Name")

parser.add_argument("-Introns", action='store_true', help="Include Intronic PASs")
args = parser.parse_args()

Root = str(args.Root)
Input_Gene = Root + '_Gene_DESeq2-results.csv'
Input_Exon = Root + '_Exon_DESeq2-results.csv'
Input_PAS = Root + '_PAS_DESeq2-results.csv'
if args.Introns:
    IncIntrons = True
else:
    IncIntrons = False

if args.Replicates:
    Replicates = int(args.Replicates)
else:
    Replicates = 3
if args.Conditions:
    Conditions = int(args.Conditions)
else:
    Conditions = 2
if args.Fraction:
    ReqFractionChange = float(args.Fraction)
else:
    ReqFractionChange = 0.1

##Read in PAS BED file into dict
PAS_library = {}
with open(str(args.PASs),'r') as In:
    line = In.readline()
    while line:
        data = line.split()
        PAS_library[data[3]] = (data[0], data[1], data[2], data[5])
        line = In.readline()

##Class for unannotated PASs
class Unannotated(object):
    def __init__(self, Name, baseMean, log2FoldChange, pvalue, padj, weight, counts):
        self.Name = Name
        self.Counts = counts
        self.baseMean = float(baseMean)
        self.Locus = PAS_library[Name]
        try:
            self.FoldChange = float(log2FoldChange)
            self.pvalue = float(pvalue)
            self.padj = float(padj)
        except:
            ##For cases were e.g. padj == 'NA'
            self.log2FoldChange = log2FoldChange
            self.pvalue = pvalue
            self.padj = padj

    def __str__(self):
        return  str([self.Name,
                self.baseMean,
                self.FoldChange,
                self.pvalue,
                self.padj])

##Class for Gene DE infor
class Gene(object):
    def __init__(self, Name, baseMean, log2FoldChange, pvalue, padj, weight, counts):
        #self.PASs = {}
        self.Exons = {}
        self.Introns = {}
        self.Name = Name
        self.Counts = counts
        self.baseMean = float(baseMean)
        try:
            self.FoldChange = float(log2FoldChange)
            self.pvalue = float(pvalue)
            self.padj = float(padj)
        except:
            self.FoldChange = log2FoldChange
            self.pvalue = pvalue
            self.padj = padj

    def FinishExons(self):
        #Conditions = 2
        #Replicates = 3
        TotalCounts = [0] * Conditions# * Replicates
        for i in self.Exons:
            TotalCounts[0] += sum(np.array(self.Exons[i].Counts[:Replicates], dtype=float))
            TotalCounts[1] += sum(np.array(self.Exons[i].Counts[Replicates:], dtype=float))
        for i in self.Exons:
            Cond1Count = sum(np.array(self.Exons[i].Counts[:Replicates], dtype=float))
            Cond2Count = sum(np.array(self.Exons[i].Counts[Replicates:], dtype=float))
            NormCond1Count = Cond1Count/TotalCounts[0]
            NormCond2Count = Cond2Count/TotalCounts[1]
            self.Exons[i].FractionChange = NormCond2Count - NormCond1Count

    def __str__(self):
        return  str([self.Name,
                self.baseMean,
                self.FoldChange,
                self.pvalue,
                self.padj,
                [i for i in self.Exons]])

##Class for Intron DE info, invoked unless IncIntron == True, then treated as exons
class Intron(object):
    def __init__(self, Name, baseMean, log2FoldChange, pvalue, padj, weight, counts):
        self.Name = Name
        self.Counts = counts
        self.PASs = {}
        self.GeneName = Name.split("_")[0]
        self.baseMean = float(baseMean)
        self.Fraction = self.baseMean / GeneDict[self.GeneName].baseMean
        self.FractionChange = 0.0
        try:
            self.FoldChange = float(log2FoldChange)
            self.pvalue = float(pvalue)
            self.padj = float(padj)
        except:
            self.FoldChange = log2FoldChange
            self.pvalue = pvalue
            self.padj = padj

    def FinishPASs(self):
        #Conditions = 2 ##Change this later or learn from metadata
        #Replicates = 3 ##Change this later or learn from metadata
        TotalCounts = [0] * Conditions# * Replicates
        for i in self.PASs:
            TotalCounts[0] += sum(np.array(self.PASs[i].Counts[:Replicates], dtype=float))
            TotalCounts[1] += sum(np.array(self.PASs[i].Counts[Replicates:], dtype=float))
        for i in self.PASs:
            Cond1Count = sum(np.array(self.PASs[i].Counts[:Replicates], dtype=float))
            Cond2Count = sum(np.array(self.PASs[i].Counts[Replicates:], dtype=float))
            NormCond1Count = Cond1Count/TotalCounts[0]
            NormCond2Count = Cond2Count/TotalCounts[1]
            self.PASs[i].FractionChange = NormCond2Count - NormCond1Count

    def __str__(self):
        return  str([self.Name,
                self.baseMean,
                self.FoldChange,
                self.pvalue,
                self.padj,
                self.Fraction,
                [i for i in self.PASs]])

##Class for Exon DE info
class Exon(object):
    def __init__(self, Name, baseMean, log2FoldChange, pvalue, padj, weight, counts):
        self.Name = Name
        self.Counts = counts
        self.PASs = {}
        self.GeneName = Name.split("_")[0]
        self.baseMean = float(baseMean)
        self.Fraction = self.baseMean / GeneDict[self.GeneName].baseMean
        self.FractionChange = 0.0
        try:
            self.FoldChange = float(log2FoldChange)
            self.pvalue = float(pvalue)
            self.padj = float(padj)
        except:
            self.FoldChange = log2FoldChange
            self.pvalue = pvalue
            self.padj = padj

    def FinishPASs(self):
        #Conditions = 2 ##Change this later or learn from metadata
        #Replicates = 3 ##Change this later or learn from metadata
        TotalCounts = [0] * Conditions# * Replicates
        for i in self.PASs:
            TotalCounts[0] += sum(np.array(self.PASs[i].Counts[:Replicates], dtype=float))
            TotalCounts[1] += sum(np.array(self.PASs[i].Counts[Replicates:], dtype=float))
        for i in self.PASs:
            Cond1Count = sum(np.array(self.PASs[i].Counts[:Replicates], dtype=float))
            Cond2Count = sum(np.array(self.PASs[i].Counts[Replicates:], dtype=float))
            NormCond1Count = Cond1Count/TotalCounts[0]
            NormCond2Count = Cond2Count/TotalCounts[1]
            self.PASs[i].FractionChange = NormCond2Count - NormCond1Count

    def __str__(self):
        return  str([self.Name,
                self.baseMean,
                self.FoldChange,
                self.pvalue,
                self.padj,
                self.Fraction,
                [i for i in self.PASs]])

##Class for annotated PASs DE info
class PAS(object):
    def __init__(self, Name, baseMean, log2FoldChange, pvalue, padj, weight, counts):
        self.Name = Name
        self.Counts = counts
        self.baseMean = float(baseMean)
        self.GeneName = Name.split("_")[0]
        self.ExonName = Name.split("_PAS")[0]
        self.Locus = PAS_library[Name]
        if 'exon' in Name or IncIntrons == True:
            self.Fraction = self.baseMean / GeneDict[self.GeneName].Exons[self.ExonName].baseMean
        else:
            self.Fraction = self.baseMean / GeneDict[self.GeneName].Introns[self.ExonName].baseMean
        self.FractionChange = 0.0
        try:
            self.FoldChange = float(log2FoldChange)
            self.pvalue = float(pvalue)
            self.padj = float(padj)
        except:
            self.log2FoldChange = log2FoldChange
            self.pvalue = pvalue
            self.padj = padj

    def __str__(self):
        return  str([self.Name,
                self.baseMean,
                self.FoldChange,
                self.padj])

GeneDict = {}
##Reads in all Gene DE info
with open(Input_Gene, 'r') as In:
        Header = In.readline()
        Headings = Header.rstrip()[1:-1].split('","')[1:]
        Data = In.readline()
        while Data:
            Data = Data.split(',')
            Name = Data[1].replace('"', '')
            GeneDict[Name] = Gene(Name, Data[2], Data[3], Data[6], Data[7], Data[8], Data[9:])
            Data = In.readline()

##Reads in all Exon DE info
with open(Input_Exon, 'r') as In:
        Header = In.readline()
        Headings = Header.rstrip()[1:-1].split('","')[1:]
        Data = In.readline()
        while Data:
            Data = Data.split(',')
            Name = Data[1].replace('"', '')
            if 'Unannotated' in Name:
                pass
            else:
                GeneName = Name.split('_')[0]
                if 'exon' in Name or IncIntrons == True:
                    GeneDict[GeneName].Exons[Name] = Exon(Name, Data[2], Data[3], Data[6], Data[7], Data[8], Data[9:])
                else:
                    GeneDict[GeneName].Introns[Name] = Intron(Name, Data[2], Data[3], Data[6], Data[7], Data[8], Data[9:])
            Data = In.readline()

##Reads in all PAS DE info
UnannotatedDict = {}
with open(Input_PAS, 'r') as In:
        Header = In.readline()
        Headings = Header.rstrip()[1:-1].split('","')[1:]
        Data = In.readline()
        while Data:
            Data = Data.split(',')
            Name = Data[1].replace('"', '')
            if 'Unannotated' in Name:
                UnannotatedDict[Name] = Unannotated(Name, Data[2], Data[3], Data[6], Data[7], Data[8], Data[9:])
            else:
                GeneName = Name.split('_')[0]
                if 'exon' in Name or IncIntrons == True:
                    ExonName = Data[1].replace('"', '').split('_PAS')[0]
                    GeneDict[GeneName].Exons[ExonName].PASs[Name] = PAS(Name, Data[2], Data[3], Data[6], Data[7], Data[8], Data[9:])
                else:
                    IntronName = Data[1].replace('"', '').split('_PAS')[0]
                    GeneDict[GeneName].Introns[IntronName].PASs[Name] = PAS(Name, Data[2], Data[3], Data[6], Data[7], Data[8], Data[9:])
            Data = In.readline()
#pickle.dump(GeneDict, open(Root + '_GeneDict.py.pi','w'))


#After reading in, calculate adbundance of PASs within genes and within exons/introns
for i in GeneDict:
    GeneDict[i].FinishExons()
    for j in GeneDict[i].Exons:
        GeneDict[i].Exons[j].FinishPASs()
    for j in GeneDict[i].Introns:
        GeneDict[i].Introns[j].FinishPASs()


def FindSizeChange(TEMP, Index, PASResult):
    Direction = TEMP[0].Locus[3]
    if Index == 0:
        if Direction == '+':
            if PASResult == 'UP':
                return "_Shortening"
            else:
                return "_Lengthening"
        else:
            if PASResult == 'UP':
                return "_Lengthening"
            else:
                return "_Shortening"
    elif Index == len(TEMP) - 1:
        if Direction == '+':
            if PASResult == 'UP':
                return "_Lengthening"
            else:
                return "_Shortening"
        else:
            if PASResult == 'UP':
                return "_Shortening"
            else:
                return "_Lengthening"
    else:
        return "_both"


##MAKE REPORT
DEn, TEn, APAn, APATEn, APAL, APAS, APAB = 0,0,0,0,0,0,0

with open(str(args.Output),'w') as Output:
    Heading = ['Gene', 'baseMean', 'FoldChange', 'padj', 'GeneResult',
               'Exons', 'baseMean', 'FractionChange', 'padj', 'ExonResult', 'TE_Result',
               'PASs', 'baseMean', 'FractionChange', 'padj', 'PASResult', 'APA_Result']
    Output.write('\t'.join(Heading))
    Output.write ('\n')

    for i in GeneDict:
        DE = False
        TE = False
        APA = False
        ##GENE RESULTS
        if GeneDict[i].padj < 0.1 and GeneDict[i].padj != 'NA':
            if GeneDict[i].FoldChange > 0.585:
                Result = 'UP'
                DE = True
                DEn += 1
            elif GeneDict[i].FoldChange < -0.585:
                Result = 'DOWN'
                DE = True
                DEn += 1
            else:
                Result = 'NC'
        else:
            Result = 'NC'
        GeneEntry = [i, str(GeneDict[i].baseMean), str(GeneDict[i].FoldChange), str(GeneDict[i].padj), Result, '']
        Output.write('\t'.join(GeneEntry))

        ##EXONS RESULTS
        TEMP = []
        for j in GeneDict[i].Exons:
            if GeneDict[i].Exons[j].Fraction > 0.05:
                TEMP.append(GeneDict[i].Exons[j])
        ##Make sure PASs are sequential
        TEMP.sort(key=lambda x: x.Name)
        NumSites = len(TEMP)
        if NumSites < 2:
            pass#ExonEntry = [TEMP[0].Name,'0','0','0','NC', 'NA', '']
        else:
            ExonEntry = [ [], [],[],[], 'NC', 'NA', '']
            for j in TEMP:
                ExonEntry[0].append(j.Name)
                ExonEntry[1].append(j.baseMean)
                ExonEntry[2].append(j.FractionChange)
                ExonEntry[3].append(j.padj)
            #Hits = [k < 0.1 for k in ExonEntry[3]]
            Hits = []
            for k in range(len(ExonEntry[0])):
                if ExonEntry[3][k] < 0.1:
                    if ExonEntry[2][k] < -ReqFractionChange or ExonEntry[2][k] > ReqFractionChange:
                        Hits.append(True)
                    else:
                        Hits.append(False)
                else:
                    Hits.append(False)
            if Hits.count(True) == 0:
                ExonEntry[-2] = 'NotSig'
            elif Hits.count(True) == 1:
                ##Single Changer
                Exon = Hits.index(True)
                if TEMP[Exon].FoldChange > 0:
                    ExonEntry[-3] = 'Exon#:' + str(Exon + 1) + "-only_UP"
                else:
                    ExonEntry[-3] = 'Exon#:' + str(Exon + 1) + "-only_Down"
                ExonEntry[-2] = 'TE'
            else:
                Result = [[],[]]
                for k in TEMP:
                    if k.padj < 0.1:
                        if k.FoldChange > 0:
                            Result[0].append(str(TEMP.index(k) + 1))
                            Result[1].append('UP')
                        else:
                            Result[0].append(str(TEMP.index(k) + 1))
                            Result[1].append('DOWN')
                        if k.FractionChange > 0.1 or k.FractionChange < -0.1:
                            ExonEntry[-2] = 'TE'
                ExonEntry[-3] = 'Exon#:' + ','.join(Result[0]) + '_' + '-'.join(Result[1])
            if ExonEntry[-2] == 'TE':
                TEn += 1
                TE = True
            Output.write('\t'.join([str(k) for k in ExonEntry]))

        ##PAS RESULTS
        for j in GeneDict[i].Exons:
                TEMP = []
                for k in GeneDict[i].Exons[j].PASs:
                    if GeneDict[i].Exons[j].PASs[k].Fraction > 0.05:
                        TEMP.append(GeneDict[i].Exons[j].PASs[k])
                ##Make sure PASs are sequential to determine lengthening vs shortening
                TEMP.sort(key=lambda x: x.Name)
                NumSites = len(TEMP)
                if NumSites == 1:
                    pass#PASEntry = [TEMP[0].Name,'0','0','0','NC', 'NA', '']
                else:
                    PASEntry = [ [], [],[],[], 'NC', 'NA', '']
                    for k in TEMP:
                        PASEntry[0].append(k.Name)
                        PASEntry[1].append(k.baseMean)
                        PASEntry[2].append(k.FractionChange)
                        PASEntry[3].append(k.padj)
                    #Hits = [k < 0.1 for k in PASEntry[3]]
                    Hits = []
                    for k in range(len(PASEntry[0])):
                        if PASEntry[3][k] < 0.1:
                            if PASEntry[2][k] < ReqFractionChange or PASEntry[2][k] > ReqFractionChange:
                                Hits.append(True)
                            else:
                                Hits.append(False)
                        else:
                            Hits.append(False)
                    if Hits.count(True) == 0:
                        PASEntry[-2] = 'NotSig'
                    elif Hits.count(True) == 1:
                        ##Single Changer
                        PAS = Hits.index(True)
                        if TEMP[PAS].FoldChange > 0:
                            PASResult = 'UP'
                            PASEntry[-3] = 'PAS#:' + str(PAS + 1) + "-only_UP"
                        else:
                            PASResult = 'DOWN'
                            PASEntry[-3] = 'PAS#:' + str(PAS + 1) + "-only_Down"
                        PASEntry[-2] = 'APA'
                        ##Position in exon determines shortening vs lengthening
                        PASEntry[-2] += FindSizeChange(TEMP, PAS, PASResult)
                    else:
                        Result = [[],[]]
                        m = 0
                        for k in TEMP:
                            if k.padj < 0.1:
                                if k.FoldChange > 0:
                                    Result[0].append(str(TEMP.index(k) + 1))
                                    PASResult = 'UP'
                                    Result[1].append('PASResult')
                                else:
                                    Result[0].append(str(TEMP.index(k) + 1))
                                    PASResult = 'DOWN'
                                    Result[1].append('PASResult')
                                if k.FractionChange > ReqFractionChange or k.FractionChange < -ReqFractionChange:
                                    PASEntry[-2] = 'APA'
                                    PASEntry[-2] += FindSizeChange(TEMP, m, PASResult)
                            m+=1
                        PASEntry[-3] = 'PAS#:' + ','.join(Result[0]) + '_' + '-'.join(Result[1])
                    if PASEntry[-2][:3] == 'APA':
                        APAn += 1
                        APA = True
                        if 'Lengthening' in PASEntry[-2]:
                            APAL += 1
                        elif 'Shortening' in PASEntry[-2]:
                            APAS += 1
                        else:
                            APAB += 1
                    Output.write('\t'.join([str(k) for k in PASEntry]))
        Output.write ('\n')
        if TE and APA:
            APATEn += 1

print "Total genes detected: %s" % len(GeneDict)
n = 0
for i in GeneDict:
    if len(GeneDict[i].Exons) > 1:
        n+=1
print "Number of genes with multiple terminal exons:%s" % n
n=0
for i in GeneDict:
    for j in GeneDict[i].Exons:
        if len(GeneDict[i].Exons[j].PASs) > 1:
            n+=1
print "Number of exons with multiple poly(A) sites: %s" % n
print "Number of differentially expressed genes (padj<0.1): %s" % DEn
print "Number of APA events: %s" % APAn
print "\t of which %s are Lengthening, %s are Shortening: and %s are both." % (APAL, APAS, APAB)
print "Number of alternative terminal exons (TE): %s" % TEn
print "Number genes with both APA and TE: %s" % APATEn
```
