## Supplementary material for "DPAC: a tool for Differential Poly(A) Site usage from poly(A)–targeted RNAseq data": DPAC software; Scripts, Manual and TestData: Extract_pA_Lens.py

```
#!/usr/bin/python
##Last Modifed 4Oct16
import gzip
import argparse
parser = argparse.ArgumentParser()
parser.add_argument("Raw_Data", help="Info File from cutadapt")
parser.add_argument("Trimmed_Data", help="Info File from cutadapt")
parser.add_argument("Output_File", help="Desired output filename")
args = parser.parse_args()

##In = open(str(args.Input_info),'r')
Output = gzip.open(str(args.Output_File), 'wb')

def FindAs(TrimmedSeq, WindowSize, Err):
        n = 0
        for i in range(len(TrimmedSeq) - WindowSize):
                Window = TrimmedSeq[n:n+WindowSize]
                As = Window.count('A')
                ##Count through Seq until disqualifying window is found
                if As >= (WindowSize-Err):
                        n+=1
                else:
                        break
        try:
                n += Window.rindex('A')
        except:
                n=0
        ##Finds position of last A in failing or completing window
        n += 1
        return n

with gzip.open(str(args.Trimmed_Data),'rb') as CutIn:
        with gzip.open(str(args.Raw_Data),'rb') as RawIn:
                ##Read Raw
                RawName = RawIn.readline()
                RawSeq = RawIn.readline().rstrip()
                RawIn.readline()
                RawIn.readline()
                ##Raw Trimmed
                Data = CutIn.readline().rstrip()
                Seq = CutIn.readline()
                DataName = CutIn.readline()
                Quals = CutIn.readline()
                ##Care that reads are in order
                while Data:
                        Name = Data.split()[0]
                        while Name not in RawName:
                        #while Name[:-16] not in RawName:
                                RawName = RawIn.readline()
                                RawSeq = RawIn.readline().rstrip()
                                RawIn.readline()
                                RawIn.readline()
                        Coord = len(Seq)# Adjust if Trimmed data + 6
                        Adaptor = RawSeq[Coord:]
                        if len(Adaptor) > 10:
                                	pA_length = FindAs(Adaptor, 10, 1)
                                #if pA_length > 15:
                                        Output.write(Name +'_pA=' + str(pA_length) + '\n')
                                        Output.write(Seq + '+\n')
                                        Output.write(Quals)
                                #else:
                                 #       pass
                        else:
                                pass
                        ##Restart
                        Data = CutIn.readline()
                        Seq = CutIn.readline()
                        DataName = CutIn.readline()
                        Quals = CutIn.readline()
Output.close()
```
