## Supplementary material for "DPAC: a tool for Differential Poly(A) Site usage from poly(A)–targeted RNAseq data": DPAC software; Scripts, Manual and TestData: Fix_BEDnames.py

```
#!/bin/python
##Last Modifed Nov18 by ALR
import argparse

parser = argparse.ArgumentParser()
parser.add_argument("Input", help="Input BED file with unclustered PASs e.g. hg19_PACs.bed. Must be sorted by count e.g. '$ sort -k4 -rn In.bed > In.sorted.bed'")
parser.add_argument("Output", help="Unmasked Output BED file for clustered annotated PASs")
parser.add_argument("Names", help="AccessionNames fasta")
args = parser.parse_args()

InFile = str(args.Input)
NameFile = str(args.Names)
Output = str(args.Output)

Names = {}
with open(NameFile, 'r') as In:
    line = In.readline()
    while line:
        Data = line.split()
        Names[Data[0]] = Data[1]
        line = In.readline()
        

AllPASs = {}
m=0
Header = ''
with open(InFile, 'r') as In:
    line = In.readline()
    while line:
        if 'track' not in line:
            Data = line.split()
            if Data[3] == '.':
                m+=1
                Exon = Data[0] + ':' + Data[1]
                Name = "Unannotated_intergenic" + "_" + Data[0] + ':' + Data[1] + "_" + "PAS-" + str(m)
            else:
                Exon = Data[3].split("_")
                n=1
                Accession = Exon[0] + "_" + Exon[1]
                Root = Names[Accession]
                Name = Root + "_" + Exon[2] + "_" + Exon[5]  + ":" + Exon[6] + "_PAS-" + str(n)
                while Name in AllPASs:
                    n+=1
                    Name = Root + "_" + Exon[2] + "_" + Exon[5]  + ":" + Exon[6]+ "_PAS-" + str(n)
            AllPASs[Name] = Data
        else:
            Header = line
        line = In.readline()

with open(Output, 'w') as Out:
    Out.write(Header)
    for i in AllPASs:
        Out.write('\t'.join([AllPASs[i][0], AllPASs[i][1], AllPASs[i][2], i,  AllPASs[i][4], AllPASs[i][5], AllPASs[i][6], AllPASs[i][7], '\n']))

                
                
```
