## Supplementary material for "DPAC: a tool for Differential Poly(A) Site usage from poly(A)–targeted RNAseq data": DPAC software; Scripts, Manual and TestData: Make_Gene-Exon-PASs_database.py

```
import argparse

parser = argparse.ArgumentParser()
parser.add_argument("Meta_Data", help="String of metadata files")
parser.add_argument("Output", help="Merged Output Root name")
args = parser.parse_args()

Inputs = str(args.Meta_Data)
Output = str(args.Output)

PASs = {}
Exons = {}
Genes = {}
Roots = []

##Compile metadata
with open(Inputs,'r') as In:
        header = In.readline()
        sample = In.readline()
        while sample:
                Root = sample.split()[0]
                Roots.append(Root)
                sample = In.readline()

##Populate Dictionary
for Root in Roots:
    Names = set()
    print Root
    with open(Root + '_PAS_overlaps.txt','r') as In:
        data = In.readline().split()
        while data:
            #Name = data[3]
            #if Name not in Names:
                #Names.add(Name)
                PAS = data[4]
                Exon = data[4].split('_PAS')[0]
                Gene = data[4].split("_")[0]
		Count = int(float(data[3]))
                ##PASs
                if PAS not in PASs:
                    PASs[PAS] = {}
                else:
                    pass
                if Root not in PASs[PAS]:
                    PASs[PAS][Root] = Count
                else:
                    PASs[PAS][Root] += Count
                ##Exons
                if Exon not in Exons:
                    Exons[Exon] = {}
                else:
                    pass
                if Root not in Exons[Exon]:
                    Exons[Exon][Root] = Count
                else:
                    Exons[Exon][Root] += Count
                ##Gene
                if Gene not in Genes:
                    Genes[Gene] = {}
                else:
                    pass
                if Root not in Genes[Gene]:
                    Genes[Gene][Root] = Count
                else:
                    Genes[Gene][Root] += Count
            	data = In.readline().split()

PAS_Events = {}
for i in PASs:
        event = i + '\t'
        temp = []
        for j in Roots:
                if j in PASs[i]:
                        temp.append(str(PASs[i][j]))
                else:
                        temp.append('0')
        counts = '\t'.join(temp)
        counts += '\n'
        PAS_Events[event] = counts
        
with open(Output + '_PAS-counts.txt','w') as Out:
        Out.write('\t')
        Out.write('\t'.join(Roots))
        Out.write('\n')
        for i in PAS_Events:
            Out.write(i)  
            Out.write(PAS_Events[i])       

Exon_Events = {}
for i in Exons:
        event = i + '\t'
        temp = []
        for j in Roots:
                if j in Exons[i]:
                        temp.append(str(Exons[i][j]))
                else:
                        temp.append('0')
        counts = '\t'.join(temp)
        counts += '\n'
        Exon_Events[event] = counts
        
with open(Output + '_Exon-counts.txt','w') as Out:
        Out.write('\t')
        Out.write('\t'.join(Roots))
        Out.write('\n')
        for i in Exon_Events:
            Out.write(i)  
            Out.write(Exon_Events[i])

Gene_Events = {}
for i in Genes:
        event = i + '\t'
        temp = []
        for j in Roots:
                if j in Genes[i]:
                        temp.append(str(Genes[i][j]))
                else:
                        temp.append('0')
        counts = '\t'.join(temp)
        counts += '\n'
        Gene_Events[event] = counts

with open(Output + '_Gene-counts.txt','w') as Out:
        Out.write('\t')
        Out.write('\t'.join(Roots))
        Out.write('\n')
        for i in Gene_Events:
            Out.write(i)  
            Out.write(Gene_Events[i])
                
```
