## Supplementary material for "DPAC: a tool for Differential Poly(A) Site usage from poly(A)–targeted RNAseq data": DPAC software; Scripts, Manual and TestData: MakeBEDGraph_pALenAr.py

```
#!/usr/bin/python

import argparse
from re import findall
import gzip
try:
        import pysam
except:
        print "Pysam not installed; no support for .BAM files."

parser = argparse.ArgumentParser()
parser.add_argument("Input_Sam", help="Sam File")
parser.add_argument("--Description", help= "Description in string format")
args = parser.parse_args()

if str(args.Input_Sam)[-6:] == 'sam.gz':
        In = gzip.open(str(args.Input_Sam),'rb')
elif str(args.Input_Sam)[-4:] == '.bam':
        In = pysam.AlignmentFile(str(args.Input_Sam),'rb')
else:
        In = open(str(args.Input_Sam),'r')

if args.Description:
        Description = str(args.Description)
else:
        Description = 'polyA'

Dict = {}   
Scoring_Scheme = {20:0,
                  21:1,
                  22:2,
                  23:4,
                  24:9,
                  25:16,
                  26:25,
                  27:36,
                  28:49,
                  29:64,
                  30:81,
                  31:100}
def Score_pA(pAs):
        n = sum(pAs)
        Score = 0
        m=21
        for i in pAs[20:31]:
                Score += i*Scoring_Scheme[m]
                m+=1
        Score += sum(pAs[31:]) * 100
        return Score

for line in In:
        Data = line.split()
        if Data[0][0] != '@':
                #Not a Header
                if Data[1] == '0' or Data[1] == '16':
                        #Only primary alignments (SE reads only)
                        if Data[1] == '0':
                                Dir = '+'
                        elif Data[1] == '16':
                                Dir = '-'
                        else:
                                pass
                        Cigar = findall(r"[^\W\d_]+|\d+", Data[5])
                        ## Breaks Cigar string in to numbers and characters
                        Seq = Data[9]
##                                Ref = 'chr' + Data[2]
                        Name = Data[0]
                        pALength = int(Name.split('_pA=')[1])
                        Ref = Data[2]
                        if Dir == '+':
                                StartPos = int(Data[3]) - 1
                                #SAM format in 1=leftmost nt
                                n=0
                                for i in Cigar:
                                        if i == 'D' or i == 'N' or i == 'M':
                                                ## Deletion, Recombination and Mapping
                                                StartPos += int(Cigar[n-1])
                                        elif i == 'I':
                                                ##Insertion in Read relative to reference
                                                StartPos -= int(Cigar[n-1])
                                        else:
                                                ##Other mappings types?
                                                ##Pads here 'S'
                                                pass
                                        n+=1
                                EndPos = StartPos + 1
                                ##Giveswidth to BEDGraph file
                        else:
                                EndPos = int(Data[3])
                                StartPos = int(Data[3]) - 1
                        DataPoint = (Ref + '\t' + str(StartPos) + '\t' + str(EndPos) + '\t' + Dir)
                        if DataPoint not in Dict:
                                Dict[DataPoint] = [1,[0]*300]
                                Dict[DataPoint][1][pALength] += 1
                        else:
                                Dict[DataPoint][0] += 1
                                Dict[DataPoint][1][pALength] += 1  
In.close()

Out= open(str(args.Input_Sam) + '.bedgraph','w')
Out.write('track type=bedGraph name="' + str(Description) + '" description="' + str(Description) + '"\talwaysZero=on\t\n')
for i in Dict:
        if i[-1] == '+':
                Out.write(i[:-1] + '\t' + str(sum(Dict[i][1])) + '\t+\t' + str(Score_pA(Dict[i][1])) + '\t' + str(Dict[i][1]) + '\t\n')
        else:
                Out.write(i[:-1] + '\t' + str(sum(Dict[i][1])) + '\t-\t' + str(Score_pA(Dict[i][1])) + '\t' + str(Dict[i][1]) + '\t\n')
Out.close()
    
```
