## Supplementary material for "DPAC: a tool for Differential Poly(A) Site usage from poly(A)–targeted RNAseq data": DPAC software; Scripts, Manual and TestData: MakeBEDGraph_pALenAr_fromMeta.py

```
#!/usr/bin/python

import argparse
import gzip
from re import findall

parser = argparse.ArgumentParser()
parser.add_argument("MetaFile", help="Meta Data File")
parser.add_argument("--Description", help= "Description in string format")
parser.add_argument("--MinScore", help= "Description in string format")
parser.add_argument("--MinCount", help= "Description in string format")

args = parser.parse_args()

if args.Description:
    Description = str(args.Description)
else:
    Description = 'PACSeq'
    
if args.MinScore:
    MinScore = int(args.MinScore)
else:
    MinScore = 1
    
if args.MinCount:
    MinCount = int(args.MinCount)
else:
    MinCount = 1

Dict = {}   
Scoring_Scheme = {20:0,
                  21:1,
                  22:2,
                  23:4,
                  24:9,
                  25:16,
                  26:25,
                  27:36,
                  28:49,
                  29:64,
                  30:81,
                  31:100}
def Score_pA(pAs):
        Score = 0
        m=21
        for i in pAs[20:31]:
                Score += i*Scoring_Scheme[m]
                m+=1
        Score += sum(pAs[31:]) * 100
        return Score
        
###MAIN SCRIPT        
In = open(str(args.MetaFile),'r')
line = In.readline()
line = In.readline()
while line:
    line = line.split()
    FileName = line[0] + '_Hisat2-mapping.sam.gz'
    print FileName
    for Data in gzip.open(FileName, 'rb'):
        Data = Data.split()
        if Data[0][0] != '@':
            #Not a Header
            if Data[1] == '0' or Data[1] == '16':
                #Only primary alignments (SE reads only)
                if Data[1] == '0':
                    Dir = '+'
                elif Data[1] == '16':
                    Dir = '-'
                else:
                    pass
                Cigar = findall(r"[^\W\d_]+|\d+", Data[5])
                ## Breaks Cigar string in to numbers and characters
                Seq = Data[9]
                Name = Data[0]
                pALength = int(Name.split('_pA=')[1])
                Ref = Data[2]
                if Dir == '+':
                    StartPos = int(Data[3]) - 1
                    #SAM format in 1=leftmost nt
                    n=0
                    for i in Cigar:
                        if i == 'D' or i == 'N' or i == 'M':
                            ## Deletion, Recombination and Mapping
                            StartPos += int(Cigar[n-1])
                        elif i == 'I':
                            ##Insertion in Read relative to reference
                            StartPos -= int(Cigar[n-1])
                        else:
                            ##Other mappings types?
                            ##Pads here 'S'
                            pass
                        n+=1
                    EndPos = StartPos + 1
                    ##Giveswidth to BEDGraph file
                else:
                    EndPos = int(Data[3])
                    StartPos = int(Data[3]) - 1
                DataPoint = (Ref + '\t' + str(StartPos) + '\t' + str(EndPos) + '\t' + Dir)
                if DataPoint not in Dict:
                    Dict[DataPoint] = [1,[0]*300]
                    Dict[DataPoint][1][pALength] += 1
                else:
                    Dict[DataPoint][0] += 1
                    Dict[DataPoint][1][pALength] += 1 
    line = In.readline()
In.close()

OutF = open(Description + '_combined-PASs.f.bedgraph','wb')
OutR = open(Description + '_combined-PASs.r.bedgraph','wb')
OutF.write('track type=bedGraph name="' + str(Description) + '" description="' + str(Description) + '"\talwaysZero=on\t\n')
OutR.write('track type=bedGraph name="' + str(Description) + '" description="' + str(Description) + '"\talwaysZero=on\t\n')
for i in Dict:
    Score = Score_pA(Dict[i][1])
    Count = sum(Dict[i][1])
    if Score >= MinScore and Count >= MinCount:
        if i[-1] == '+':
            OutF.write(i[:-1] + str(Count) + '\t+\t' + str(Score) + '\t\n')
        else:
            OutR.write(i[:-1] + str(Count) + '\t-\t' + str(Score) + '\t\n')
OutF.close()
OutR.close()
    
```
