## Supplementary material for "DPAC: a tool for Differential Poly(A) Site usage from poly(A)–targeted RNAseq data": DPAC software; Scripts, Manual and TestData: PAS_Clustering.py

```
#!/bin/python
##Last Modifed Nov18 by ALR
import os
from subprocess import check_output
import argparse
import numpy as np

parser = argparse.ArgumentParser()
parser.add_argument("Input", help="Input BED file with unclustered PASs e.g. hg19_PACs.bed. Must be sorted by count e.g. '$ sort -k4 -rn In.bed > In.sorted.bed'")
parser.add_argument("Output", help="Unmasked Output BED file for clustered annotated PASs")
parser.add_argument("--Window", help="clustering window. Default = 20")
parser.add_argument("Strand", help="State Strand, F or R")
parser.add_argument("Genome", help="Genome_Path fasta")
#parser.add_argument("Names", help="AccessionNames fasta")
#parser.add_argument("Annotations", help="Genome_Path fasta")
parser.add_argument("--ACount", help="Allowed ACount. Default = 12")
args = parser.parse_args()

InFile = str(args.Input)
Genome = str(args.Genome)
#Annotations = str(args.Annotations)

if str(args.Strand) == 'F':
    Strand = '+'
elif str(args.Strand) == 'R':
    Strand = '-'
else:
    print "ERROR: incorrect strandedness entered, must be 'F' of 'R'."
    Strand = 'STRAND'
    
if args.ACount:
    ACount = int(args.ACount)
else:
    ACount = 12
    
if args.Window:
    Window = int(args.Window)
else:
    Window = 25    

###################
def Rev_Comp(Seq):
        Seq = Seq.upper()
        basecomplement = {'A': 'T', 'C': 'G', 'G': 'C', 'T': 'A', 'N': 'N'}
        letters = list(Seq)
        letters = [basecomplement[base] for base in letters]
        return ''.join(letters)[::-1]

##Clustering SCRIPT
Clusters = {} #chrs[coord]: [Count, start, stop]
with open(InFile, 'r') as In:
    Data = In.readline()
    while Data:
        if 'track' not in Data:
            Data = Data.split('\t')
            Chr = Data[0]
            PAS = int(Data[1])
            Count = int(Data[3])
            if Chr not in Clusters:
                Clusters[Chr] = {}
                Clusters[Chr][PAS] = [Count, PAS, PAS+1]
            else:
                NewCluster = True
                for i in Clusters[Chr]:
                    if PAS > Clusters[Chr][i][1] - Window and PAS < Clusters[Chr][i][2] + Window:
                        Clusters[Chr][i][0] += Count
                        Clusters[Chr][i][1] = min(PAS, Clusters[Chr][i][1])
                        Clusters[Chr][i][2] = max(PAS, Clusters[Chr][i][2])
                        NewCluster = False
                        break
                    else:
                        pass#new cluster
                if NewCluster:
                    Clusters[Chr][PAS] = [Count, PAS, PAS+1]
        else:
            print 'Track is:',  Data
        Data = In.readline()

Output = open('UNMASKED_' + str(args.Output), 'w')
#x = Header.index('name') + 6
#NewHeader = 'track name="Clustered_' + Header[x:]
#Output.write(NewHeader)

NAME = 'Unmasked'
for i in Clusters:
    for j in Clusters[i]:
        line = i, str(Clusters[i][j][1]), str(Clusters[i][j][2]), NAME, str(Clusters[i][j][0]), Strand, str(j), str(j+1), 
        line = '\t'.join(line)
        Output.write(line + '\n')
Output.close()

Output = open('UNNAMED_' + str(args.Output), 'w')
with open('UNMASKED_' + str(args.Output), 'r') as In:
        line = In.readline()
        while line:
                Data = line.split('\t')
                if Strand == "+":
                        cmd = Data[0] + ":" + Data[6] + "-" + str(int(Data[6]) + 20)
                elif Strand == "-":
                        cmd = Data[0] + ":" + str(int(Data[6]) - 20) + "-" + Data[6]
                else:
                        print "Enter either F or R for strandedness"
                        break
                Seq = check_output(['samtools', 'faidx', Genome, cmd])
                try:
                        Seq = Seq.split()[1]
                except:
                        cmd = 'chr' + cmd
                        line = 'chr' + line
                        Seq = check_output(['samtools', 'faidx', Genome, cmd])
                        try:
                                Seq = Seq.split()[1]
                        except:
                                Seq = ''
                if Seq:
                        Seq = Seq.upper()
                        if Strand == "-":
                                Seq = Rev_Comp(Seq)
                        else:
                                pass
                        Count = [Seq.count("A"), Seq.count("T"), Seq.count("G"), Seq.count("C")]

                        if Count[0] > ACount:
                                pass
                        else:
                               Output.write(line)
                else:
                        print "Mask_ints.py:  Failed locus in index: ", cmd[3:]
                line = In.readline()
Output.close()
#print "For both F and R strands, next command is: 'intersectBed -wao -s -a UNNAMED_" + str(args.Output), "-b <Annotations> | sort -u -k1,1 -k2,2n | awk '{OFS=\"\t\"}{print $1, $2, $3, $12, $5, $6, $7, $8}' >",  str(args.Output)
```
